# Transcriptional analysis identifies novel biomarkers associated with successful ex-vivo perfusion of human donor lungs

**DOI:** 10.1101/612374

**Authors:** John R. Ferdinand, Morvern I. Morrison, Anders Andreasson, Catriona Charlton, Alisha Chhatwal, William E. Scott, Lee A. Borthwick, Menna R. Clatworthy, Andrew J. Fisher

## Abstract

Transplantation is an effective treatment for end-stage lung disease but donor organ shortage is a major problem. *Ex-vivo* lung perfusion (EVLP) of marginal organs enables functional assessment under normothermic conditions to facilitate clinical decision-making around utilisation, but the molecular processes occurring during EVLP, and how they differ between more or less viable lungs, remains to be determined. Here we used RNA sequencing to delineate changes in gene expression occurring in n=10 donor lungs undergoing EVLP, comparing lungs that were deemed transplantable (n=6) to those deemed unusable (n=4). We found that lungs deemed suitable for transplantation following EVLP had reduced induction of a number of innate immune pathways during EVLP, but a greater increase in genes involved in oxidative phosphorylation, a critical ATP-degenerating pathway. Furthermore, *SCGB1A1*, a gene encoding an anti-inflammatory secretoglobin CC10, and other club cell genes were significantly increased in transplantable lungs following perfusion, whilst *CHIT-1* was decreased. Using a larger validation cohort (n=18), we confirmed that the ratio of CHIT1 and SCGB1A1 protein levels in lung perfusate have potential utility to distinguish transplantable and non-transplantable lungs (AUC 0.81). Together, our data identify novel biomarkers that may assist with pre-transplant lung assessment, as well as pathways that may amenable to therapeutic intervention during EVLP.

**Single sentence summary:** Transcriptional changes in lungs undergoing *ex vivo* normothermic perfusion identify chitinase1 and club cell genes as potential biomarkers to guide utilisation

## Introduction

Lung transplantation is an effective treatment for selected patients with life-threatening end-stage lung disease. Unfortunately, a shortage of donor organs limits access to transplantation for many patients who might benefit from this treatment, some of whom do not survive the wait for a suitable organ (1). One strategy to address the challenge of organ shortage is to increase the donor organ pool by utilising lungs retrieved from extended criteria donors (ECD) (1, 2). These donors do not fully satisfy the characteristics of an ideal lung donor due to advanced age, smoking history, lung function or co-morbidity (3–8) and potentially have an increased risk of poor outcomes if sub-optimal organs are inadvertently used (9, 10). A potential solution to this problem is to use ex-vivo lung perfusion (EVLP) to gather information on the organ’s function to facilitate decision-making around utilisation (11–13).

EVLP has emerged over the last 10 years as a promising technique to objectively assess the function of, and potentially recondition, lungs deemed unsuitable for immediate transplantation; with the overall aim of increasing the available donor lung pool (14). The decision to accept organs for transplantation after EVLP is currently based on physiological parameters such as oxygenation, lung compliance, pulmonary vascular resistance and peak airway pressure in addition to visual appearance of the organ. Reported discard rates of perfused lungs are highly variable, ranging from 10–60%, suggesting that some donor lungs are being inappropriately used and some unnecessarily declined for transplant after EVLP (15).

The cellular and molecular events occurring during EVLP have not been well characterised, and how these processes differ between lungs that are deemed useable compared with those that are discarded is unknown. In addition, there is a pressing need to identify novel predictive biomarkers that can be utilised during EVLP. Cellular injury occurring during the process of lung retrieval and storage results in the release of danger signals that activate immune cells, so-called sterile inflammation, causing further collateral tissue damage (16, 17). We previously demonstrated the feasibility of measuring pro-inflammatory and tissue injury signals in perfusion fluid during EVLP to identify unsuitable donor lungs (18). Furthermore, we recently demonstrated that interleukin (IL)-1β and tumour necrosis factor (TNF) levels in perfusate after 30 minutes of EVLP are predictive of in-hospital mortality post-transplantation (19). Although these candidate proteins in perfusate may have potential utility as biomarkers, their selection is based on published literature that identified proteins associated with primary lung graft dysfunction (20–25) and does not provide a comprehensive means of assessing the molecular pathways impacted by EVLP nor of identifying novel biomarkers.

Here we utilized RNA-sequencing to generate an unbiased profile of the transcriptome of human donor lungs prior to, and following EVLP, comparing changes in gene expression between lungs deemed suitable for transplantation, and those discarded on the basis of standard physiological parameters. During EVLP, we observed an increase in a number of immune pathway genes. When comparing transplantable versus discarded lungs, we found that immune pathway genes, including NLRP3 inflammasome-associated genes, positively correlated with lung discard as did *CHIT1* (encoding chitotriosidase), whilst genes involved in energy generation by oxidative phosphorylation (OXPHOS) negatively correlated with lung discard. Furthermore, *SCGB1A1*, an anti-inflammatory secretoglobin, and other club cell genes, were also significantly increased in transplantable lungs. Overall, our data suggest that lungs deemed suitable for transplantation following EVLP have reduced induction of multiple immune pathways, are better able to generate ATP and may have increased club cell number or activation than those that are discarded. Our study identifies specific outcome-associated biomarkers and pathways amenable to therapeutic intervention during EVLP that will inform the design of future clinical trials in this area.

## Results

### Transcriptomic analysis of lungs pre and post EVLP reveals an induction of immune pathway genes

Between April 2012 and July 2014, 53 pairs of human donor lungs deemed unsuitable for immediate transplantation underwent EVLP as part of the DEVELOP-UK clinical trial (26). Pre- and post-EVLP (Lund protocol, see methods) lung tissue samples were available from n=10 donor lungs and underwent RNA sequencing. Of these, EVLP was deemed successful (“pass”) in n=6 and unsuccessful (“fail”) in n=4 using standard donor lung assessment criteria (26). The donor demographics and perfusion characteristics of these lungs are described in **Table 1**.

**Table1:** Sample information

| **Parameter** | **Median (IQR)** |
| --- | --- |
| **Age** | 51 (46.75-52.75) |
| **Sex** | 7 Male 3 Female |
| **Outcome of EVLP** | 6 Pass 4 Fail |
| **Donor TLC (L)** | 6.42 (5.46-6.94) |
| **Donor type** | 2 DCD 8 DBD |
| **Smoker** | 4 Yes 6 No |
| **Ventilation prior to organ retrieval (days)** | 2.15 (1.45-3.63) |
| **Cold Ischemic time (min)** | 218 (200.5-242.25) |

We first sought to investigate changes in gene expression occurring in lungs during EVLP by comparing pre- and post-EVLP samples (**Fig. 1A**). We found that 293 genes were significantly up-regulated, and 78 genes significantly down-regulated during EVLP (**Fig. 1B, C**). Several of the top 20 most significantly upregulated genes were known to be involved in immune processes including *CD83* and interferon regulatory factor (IRF)1 (**Fig. 1C**). IRF1 is a transcription factor that drives the expression of a number of immune genes, including interferon(IFN)ß (27), a cytokine with some anti-inflammatory effects. We also observed an increase in other transcription factors known to play a central role in controlling immune cell activation, including *NFKB* (**Fig. S1A**). Consistent with this, *NFKBIE*, a negative regulator of NFkB pathways was also significantly up-regulated during EVLP (**Fig. 1C**). Heat shock proteins (HSPs) are protein chaperones that contribute to the canonical protective cellular response to a variety of physiological stressors (28) but also act as innate immune stimulators when released extracellularly (29). The HSP70 family proteins *HSPA1A* and *HSPH1* (also known HSP105) were among the genes most significantly upregulated during EVLP (**Fig. 1C**).

**Figure 1.**
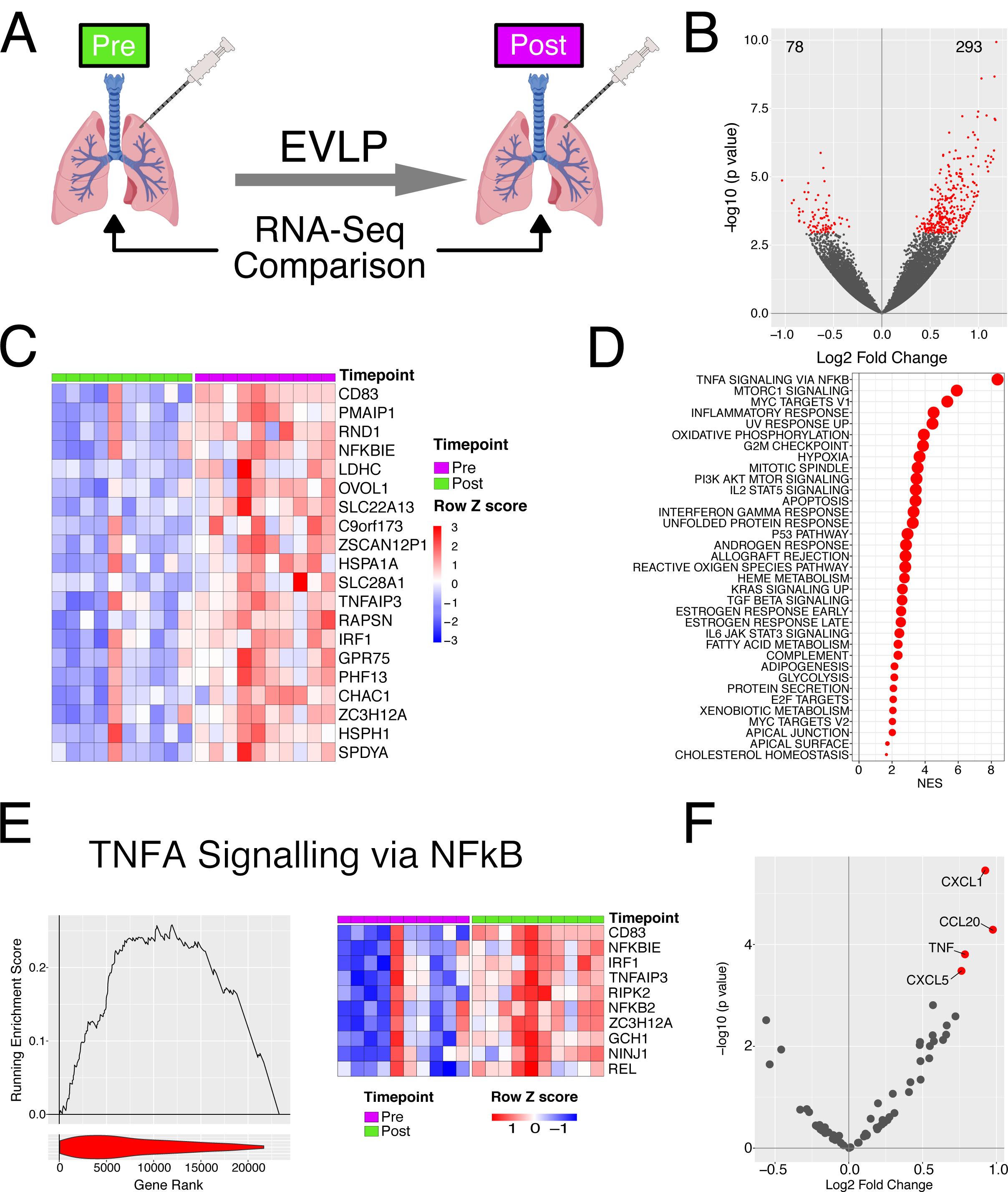
Transcriptional changes occur during EVLP. **A**-Diagrammatic representation of the experiment. **B** - Volcano plot comparing transcriptome pre and post perfusion. Red points indicate significantly differentially expressed genes (BH adjusted p value < 0.05 and are enumerated on the plot. **C** – Heatmap of top 20 significant genes upregulated during EVLP. **D** – GSEA analysis against the Hallmarks dataset for EVLP. All significantly enriched pathways (FDR q value < 0.05) have been plotted, size of point is inversely correlated to the FDR q value, red points indicated positively enriched pathways and blue negatively enriched. **E** – Left panel. GSEA enrichment for the TNF via NFkB pathway. Line plot indicates the running enrichment score and violin plot the distribution of genes from the pathways of interest within the ranked gene list. Right panel - top 10 ranked genes within the leading edge genes from the TNFa via NFkB pathway. **F** – Volcano plot comparing transcriptome pre and post perfusion showing only cytokine and chemokine genes. Red points indicate significantly differentially expressed genes (BH adjusted p value < 0.05).

We next assessed changes in molecular pathways rather than individual genes. This gains statistical power by considering the expression of groups of genes within a common pathway rather than individual genes. We performed gene set enrichment analysis (GSEA) using gene-sets curated in the ‘Hallmarks’ database. This confirmed immune system activation during EVLP, identifying ‘TNFA signalling via NFkB’ pathway as the most enriched in the dataset, and enrichment of other immune pathway genes including ‘Inflammatory response’, ‘IL2-STAT5 signalling’, and ‘Interferon gamma response’ pathways (**Fig. 1D, E**). Notably, a number of metabolic pathways were also enriched including ‘Oxidative phosphorylation’ and ‘Fatty acid metabolism’ (**Fig. 1D**), both of which contribute to the generation of adenosine triphosphate (ATP), a critical energy source for cells. Together, these data show that during EVLP, a number of immune pathways are induced as well as pathways involved in energy generation.

To further characterise immune activation, we assessed chemokine and cytokine genes, and found a significant increase in the expression of four genes during EVLP, *TNF, CXCL1* and *CXCL5* (neutrophil-recruiting chemokines), and *CCL20* (a chemokine for CCR6-expressing lymphocytes, including CD4 T cells and innate lymphoid cells) (**Fig. 1F**). We also measured a panel of cytokines in the perfusates obtained from the EVLP circuit of the donor lungs included in our transcriptomic study to assess RNA-protein correlates. This demonstrated a significant positive correlation between transcript expression in pre-EVLP biopsies and cytokine concentration in the post-EVLP perfusate sample for CCL2, CXCL8, IL8 and ICAM1 (**Fig. S1B**). In contrast, post-EVLP transcript levels showed no significant correlation with any perfusate cytokine (**Fig. S1C**). This suggests that proteins released during perfusion reflect pathways already activated in the tissue prior to EVLP.

### Lungs deemed unusable have increased induction of immune pathway genes following EVLP compared with those deemed transplantable

One of the aims of EVLP is to assess organ function in order to determine suitability for transplantation. In the n=10 lungs studied, n=6 were deemed transplantable (”Pass”) and n=4 were deemed unusable (“Fail”) based on standard donor lung assessment criteria (**Fig. 2A**). We therefore sought to determine if there were differences in the tissue transcriptome between transplantable and unusable organs, either prior to, during, or after EVLP. We found no significant differences in gene expression in pre-EVLP samples when comparing Pass and Fail lungs (data not shown). In post-EVLP samples, 12 genes were significantly upregulated in unusable lungs, including genes of immunological importance, specifically *CHIT1* (encoding chitotriosidase), *CHI3L1* (encoding chitinase-3-like 1), *IL12RB2, CXCR6, SLAMF7, SLC7A5*, and *GBP5* (**Fig. 2B**). Chitotriosidase is a hydrolase produced by alveolar macrophages in response to infection (30–32) and its expression has been associated with both granulomatous and fibrotic lung diseases (33). Indeed, serum CHIT1 has been investigated as a biomarker of disease severity and progression in several lung diseases, including chronic obstructive pulmonary disease (34). Chitinase-like proteins (including CHI3L1) are expressed by alveolar macrophages and play a role in lung inflammation, remodeling and fibrosis (35, 36). Of note, SLC7A5 (an amino acid transporter) and GBP5 (Guanylate-binding protein 5) promote NLRP3 inflammasome activation and IL1ß production in monocytes and macrophages (37, 38). Taken together, these upregulated genes suggest a marked activation of alveolar macrophages in unusable lungs, likely in response to tissue damage signals.

**Figure 2.**
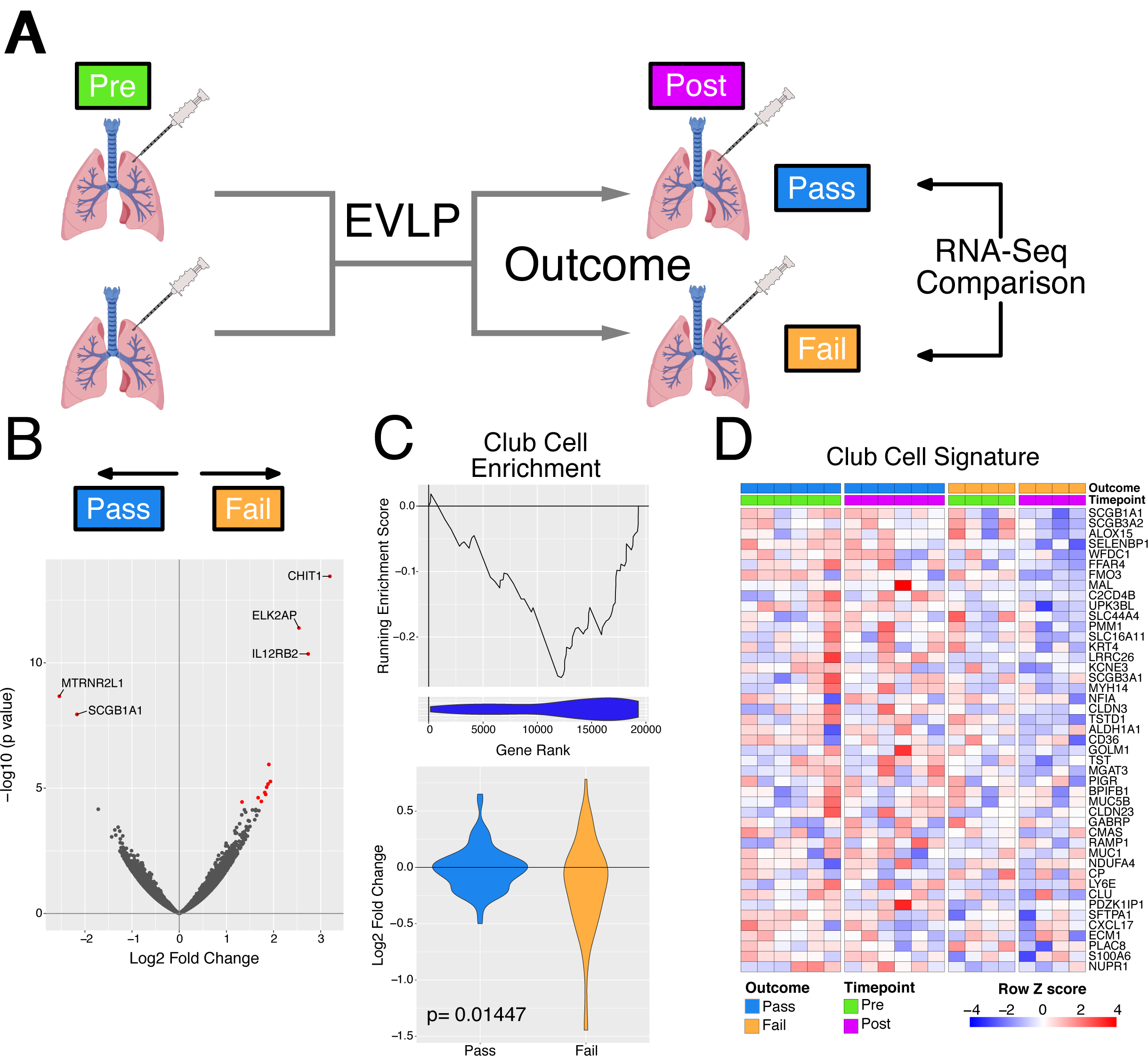
Assessment of differences in the transcriptome following EVLP with respect to outcome. **A –** Diagrammatic representation of the analysis. **B -** Volcano plot indicating results of differential expression analysis between those lungs which passed and those that failed EVLP post perfusion relative to pass. Red points indicate significantly differentially expressed genes (BH adjusted p value < 0.05). **C** – Top panel - GSEA enrichment plot for club cell marker genes from analysis in B. Line plot indicates the running enrichment score and violin plot the distribution of Club cell marker genes within the ranked gene list. Bottom panel – Violin plot indicated the log fold change in expression for all the Club cell marker genes. The p value is for comparison of the two groups using a Mann-Whitney test. Blue is organs which have passed EVLP and orange genes which have failed. **D** – Heatmap of leading edge club cell marker genes from B.

Two genes were significantly upregulated in transplantable lungs, *MTRNR2L1*, (encoding humanin-like 1), and *SCGB1A1*, a secretoglobin expressed by non-ciliated bronchiolar epithelial cells and a marker of club (previously known as ‘Clara’) cells (39) (**Fig. 2B**). Humanin proteins have been shown to protect cells from oxidative stress, starvation and hypoxia (40, 41), therefore, the induction of *MTRNR2L1* may be functionally important in donor lungs. Club cells are non-ciliated bronchiolar exocrine cells which protect the bronchiolar epithelium by producing SCGB1A1 (an anti-inflammatory secretoglobin, also known as club cell protein 10 (CC10), (42) (43) protease inhibitors, and by detoxifying harmful molecules entering the small airways via their cytochrome p450 enzymes. To further explore their presence in transplantable lungs we used a previously published single cell RNA-seq analysis of lungs to define a transcriptomic signature for club cells (44) and used this to assess enrichment for genes associated with them in perfusion. This demonstrated a significant negative enrichment of club cell genes in discarded (fail) versus transplantable (pass) lungs following perfusion (NES = −2.69, FDR q value = 0), with a greater decrease in the downregulation of these during perfusion (p=0.01, **Fig. 2C-D, S2**). We also found an enrichment of a ciliated epithelial cell and neuroendocrine cell gene signature in transplantable lungs (**Fig. S2**).

To further investigate changes in molecular pathways that differed during EVLP in transplantable versus discarded lungs, we used GSEA to compare differences between pre- and post-perfusion gene expression in pass and fail lungs (**Fig. 3A**). Positively enriched pathways that showed a greater increase during perfusion in unusable lungs were dominated by immunological processes, including ‘Allograft rejection’ and ‘IFN gamma response’ whilst the most negatively enriched pathway, ‘Oxidative phosphorylation’, showed minimal change during perfusion in those lungs which failed but an increase in those lungs which passed (**Fig. 3B-E**).

**Figure 3.**
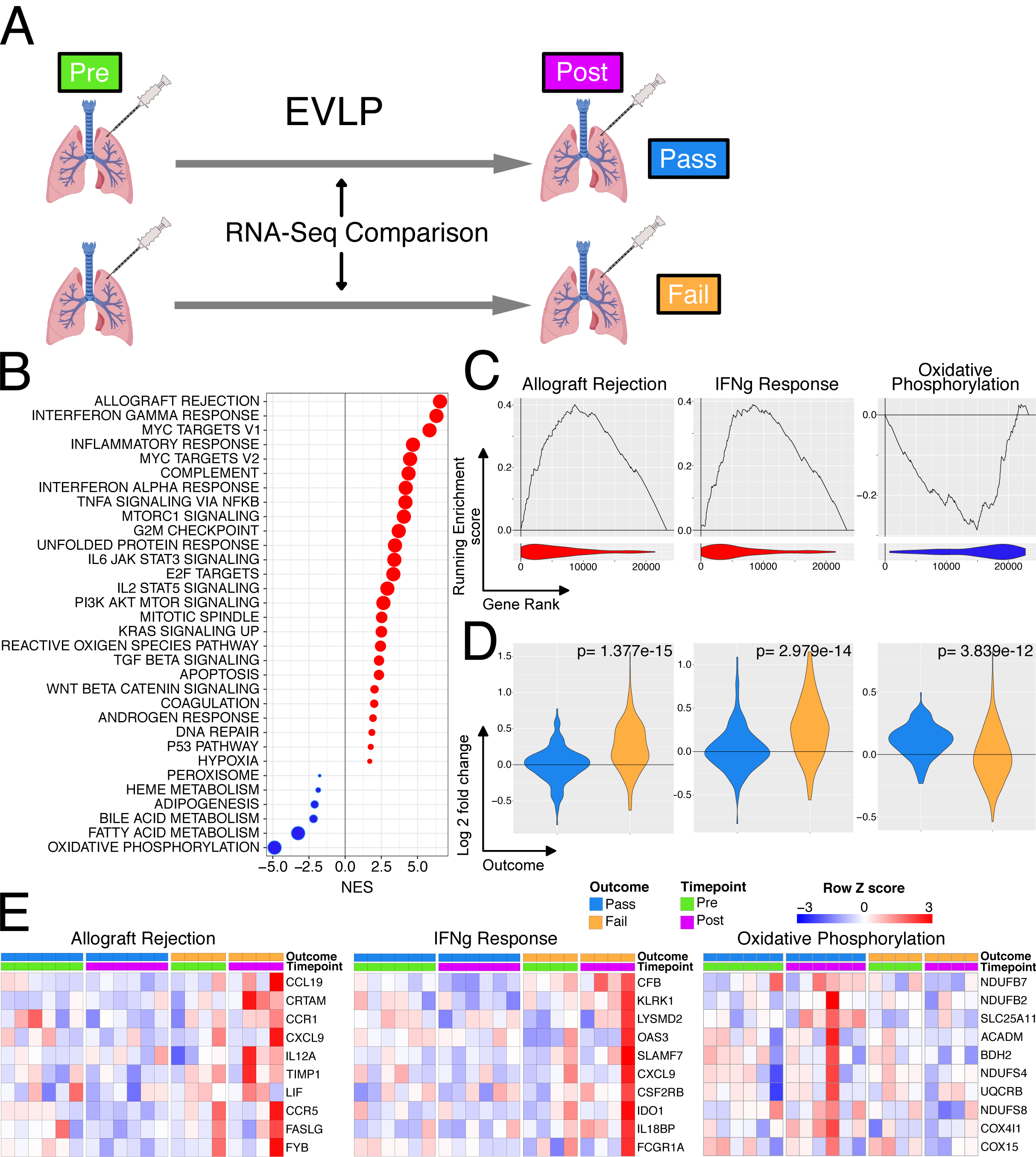
Differential gene expression during EVLP comparing lungs which pass or fail. **A –** Diagrammatic representation of the analysis. **B** - The interaction of perfusion and outcome was compared and the resulting differential expression data was used to rank genes for GSEA analysis against the Hallmarks database. All significantly enriched pathways (FDR q value < 0.05) have been plotted, size of point is inversely correlated to the FDR q value, red points indicated positively enriched pathways and blue negatively enriched. **C** - Individual enrichment plots from analysis in A. Line plot indicates the running enrichment score and violin plot the distribution of genes from the pathways of interest within the ranked gene list. **D** - Transcriptional changes were assessed for pre and post EVLP separately for those organs which passed EVLP verses those that failed. These results were filtered for only the genes in the indicated pathway above each plot and violin plots of log2 fold changes for each gene were plotted. Blue is organs which have passed EVLP and orange genes which have failed. The p value is for comparison of the two groups using a Mann-Whitney test. **E** – Top 10 genes by rank of leading edge genes for the indicated GSEA pathway when comparing the interaction of outcome and perfusion stage.

### Increase in NLRP3 inflammasome-associated genes in lungs deemed unusable

Perfusate IL-1β has been investigated as a predictive biomarker of successful EVLP and post-transplant outcome in ECD lungs, with increased perfusate IL-1β noted in donor lungs that were unusable compared with those deemed transplantable (19). The NLRP3 inflammasome acts as a major hub to generate IL-1β in response to damage-associated molecular patterns (DAMPs) (45, 46). Inflammasome assembly is a two-step process; ‘Signal 1’ is required for NLRP3 transcription, which is driven by a number of transcription factors, including NFkB, and ‘Signal 2’, including stimuli such as ATP (acting via the P2X7R), for the generation of a multimeric complex containing NLRP3, ASC and pro-caspase-1 (46). This, in turn, produces caspase-1 which cleaves pro-IL1β and pro-IL18 to their active from for secretion. A number of genes associated with inflammasome activation were increased following perfusion, particularly in lungs which failed EVLP compared to those which were deemed transplantable (**Fig. 4A**) and some members of the gene set were upregulated to a greater extent during perfusion in the organs deemed unusable (**Fig. 4B**). Overall, these data suggest that NLRP3 inflammasome activation occurs both prior to and during EVLP, and that this process is more marked in lungs that are deemed unusable after EVLP.

**Figure 4.**
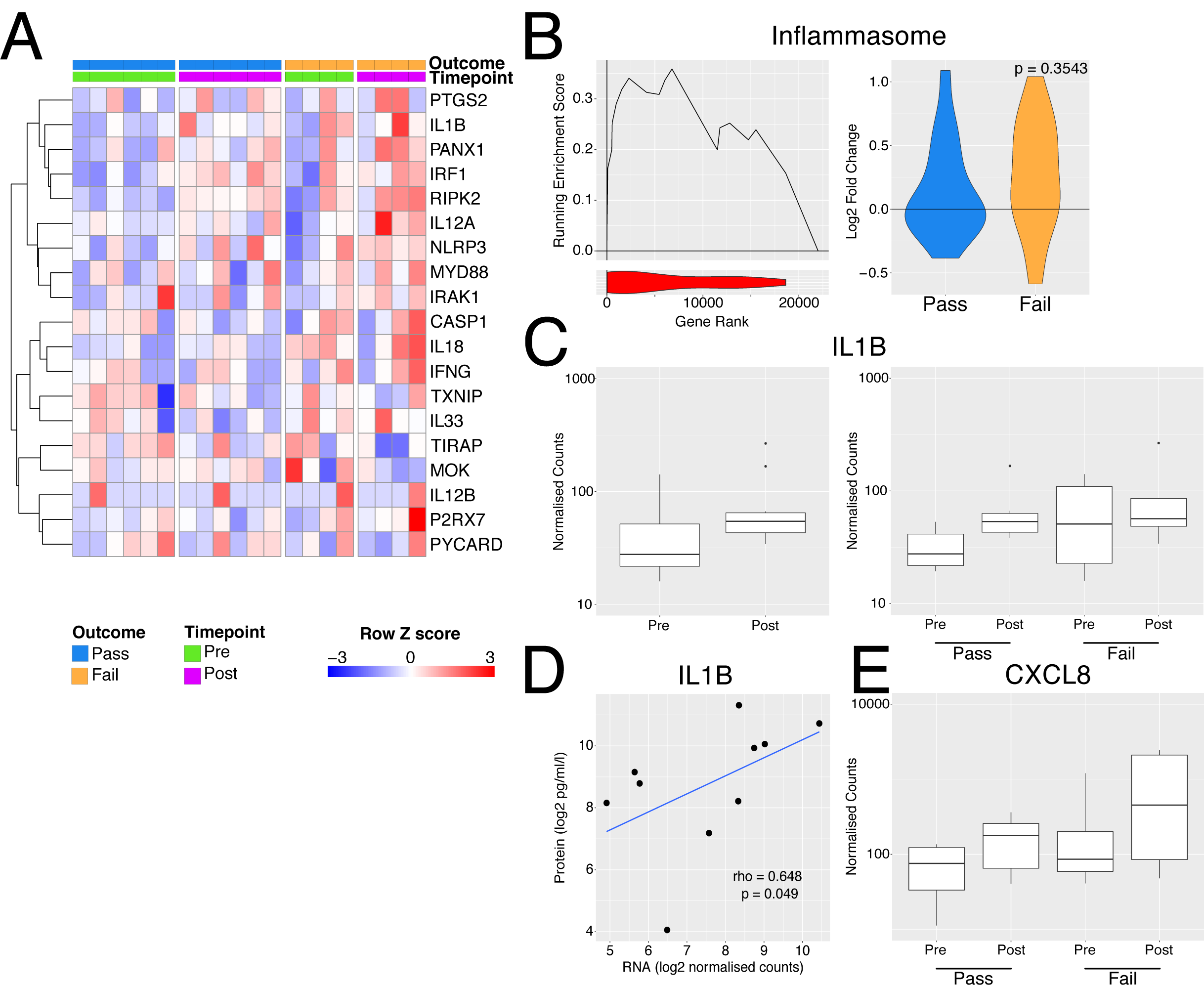
Inflammasome activation with EVLP. **A** - Heatmap of genes involved in inflammasome activation. **B** – Left panel - GSEA enrichment plot for inflammasome genes from A comparing organs which passed vs those that failed EVLP prior to undergoing perfusion. Right panel – violin plot indicated log2 fold change in inflammasome genes during perfusion separately for the pass (blue) and fail (yellow) groups. The p value is for comparison of the two groups using a Mann-Whitney test. **C** - Boxplot of normalised counts for IL1B separated by timepoint alone (left) or both timepoint and outcome (right). **D** - Scatter plot of normalised transcript expression prior to EVNP against IL1B protein in the perfusate at the end of EVLP. Correlation coefficient and p values was calculated using Spearmans test prior to log2 transformation, blue line indicates a liner model fitted to the data post transformation. **E** - Boxplot of normalised counts for CXCL8 in samples pre and post perfusion separated by outcome.

In terms of IL1 β, when considering all 10 lungs analysed, we found a trend towards an increase in transcripts in post-EVLP samples compared with those obtained pre-EVLP (**Fig. 4C**), with unusable lungs showing variable and non-significantly higher *IL1B* transcripts pre-EVLP (**Fig. 4C**). *IL1B* transcript levels in pre- (**Fig. 4D**) but not post-EVLP (data not shown) biopsies significantly correlate though weekly with the protein concentration in the perfusate, suggesting that either the RNA had not yet been translated into pro-IL1□ protein or that pro-IL1□ had not been processed into its mature form and secreted within the time-window of this experiment, likely the latter, since post-translational regulation is known to be important for IL1□ (47).

One effect of IL1β is to stimulate the production of neutrophil-recruiting chemokines, including CXCL8. There was a trend towards an increase in *CXCL8* transcripts following EVLP, particularly in discarded lungs (**Fig. 4E**) and variable increases in other neutrophil-attracting chemokines and adhesion molecules (**Fig. S3**).

### Differing patterns of expression of heat shock protein family members during EVLP

Since some HSP70 genes were among those most significantly upregulated during EVLP (**Fig. 1C**), we assessed whether there were differences in the expression of all HSP gene family members in lungs deemed transplantable and those deemed unusable. This analysis showed three patterns of gene expression: 1) Some HSP family members (gene group 1) were already induced in pre-EVLP biopsies in both Pass and Fail lungs. These showed little increase in post-EVLP biopsies in Pass lungs but did increase during EVLP in Fail lungs (**Fig. 5A, B**); suggesting that these may have deleterious effects. 2) Other HSPs (gene group 2) showed low expression in pre-EVLP biopsies in both Pass and Fail lungs, but their expression increased in post-EVLP biopsies in both Pass and Fail lungs (**Fig. 5A, B**). 3) A third group of HSP genes (gene group 3) were already expressed in pre-EVLP biopsies in both Pass and Fail lungs and their expression was maintained in post-EVLP biopsies in Pass lungs but decreased in Fail lungs (**Fig. 5A, B**); suggesting that these may have protective effects. Group 3 HSPs were predominantly HSPB genes, a family of small HSPs.

**Figure 5.**
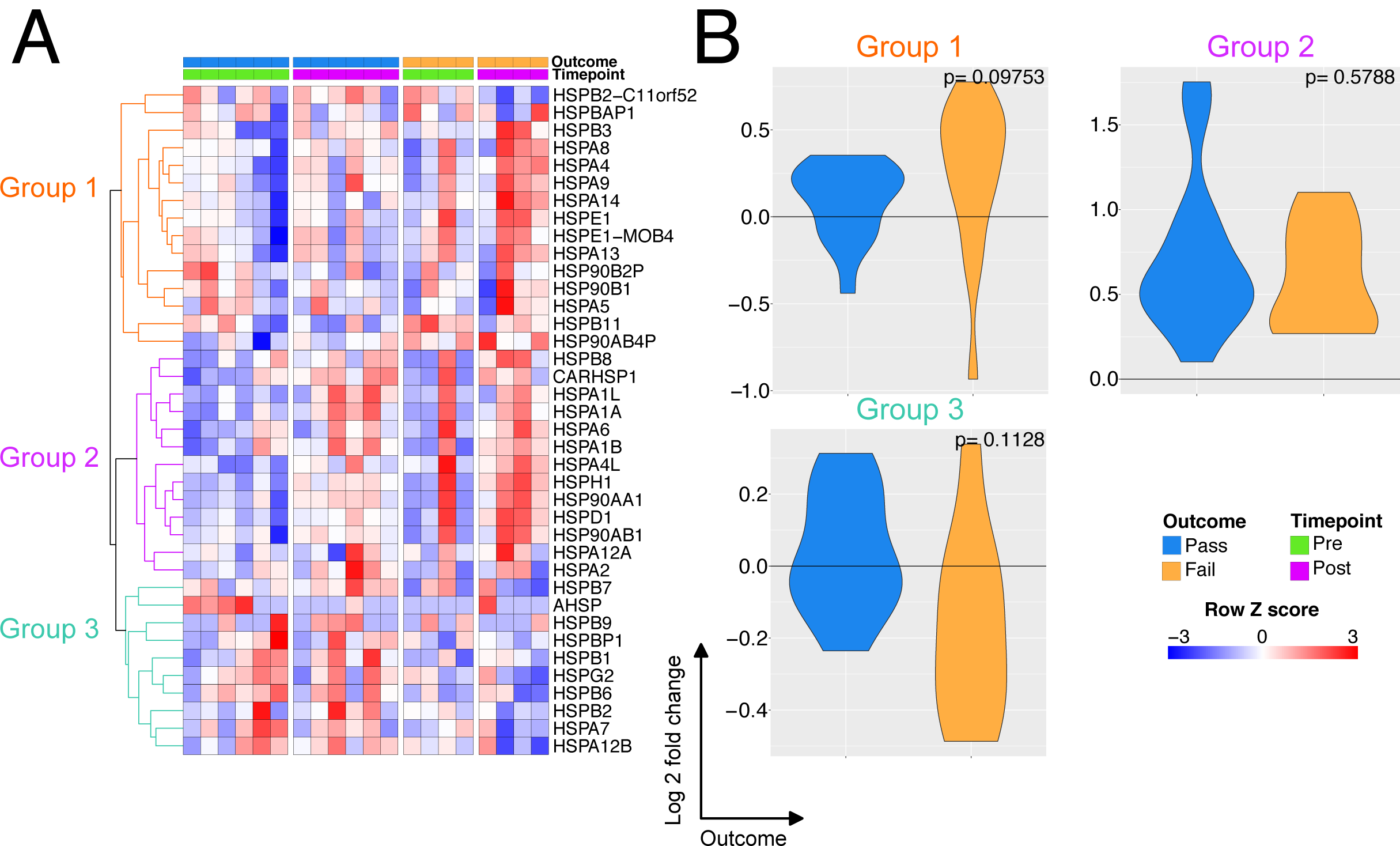
Effect of perfusion on the expression of heat shock protein transcripts. **A** - Heatmap of all heat shock protein transcripts in perfusion. The expression profiles were hierarchically clustered and 3 groups identified using a k-means approach. **B** - Transcriptional changes were assessed for pre and post EVLP separately for those organs which passed EVLP vs those that failed for each group of HSP genes. Blue is organs which have passed EVLP and orange genes which have failed. The p value is for comparison of the two groups using a Mann-Whitney test.

### Assessment of RNA and protein correlates for potential biomarkers

To explore whether the differentially expressed genes we had identified in ‘Pass’ and ‘Fail’ lungs (**Fig. 2A**) might be useful protein biomarkers of successful perfusion, we measured SCGB1A1 and chitinase 1 (CHIT1) in perfusion fluid. These were chosen because they were significantly differentially expressed genes in ‘Pass’ and ‘Fail’ lungs respectively, encoded secreted proteins, and quantification reagents/assays were commercially available. In perfusate samples taken after 150 min of perfusion with the Lund protocol in n=17 lungs taken from the DEVELOP UK study, we confirmed an increase in SCGB1A1 in transplanted lungs and a decrease in CHIT1 (**Fig. 6A**). The ratio of the concentration of both proteins were applied to a receiver operating characteristic (ROC) analysis and demonstrated reasonable positive and negative predictive value for lung utilisation (Area under the curve = 0.8, **Fig. 6B**).

**Figure 6.**
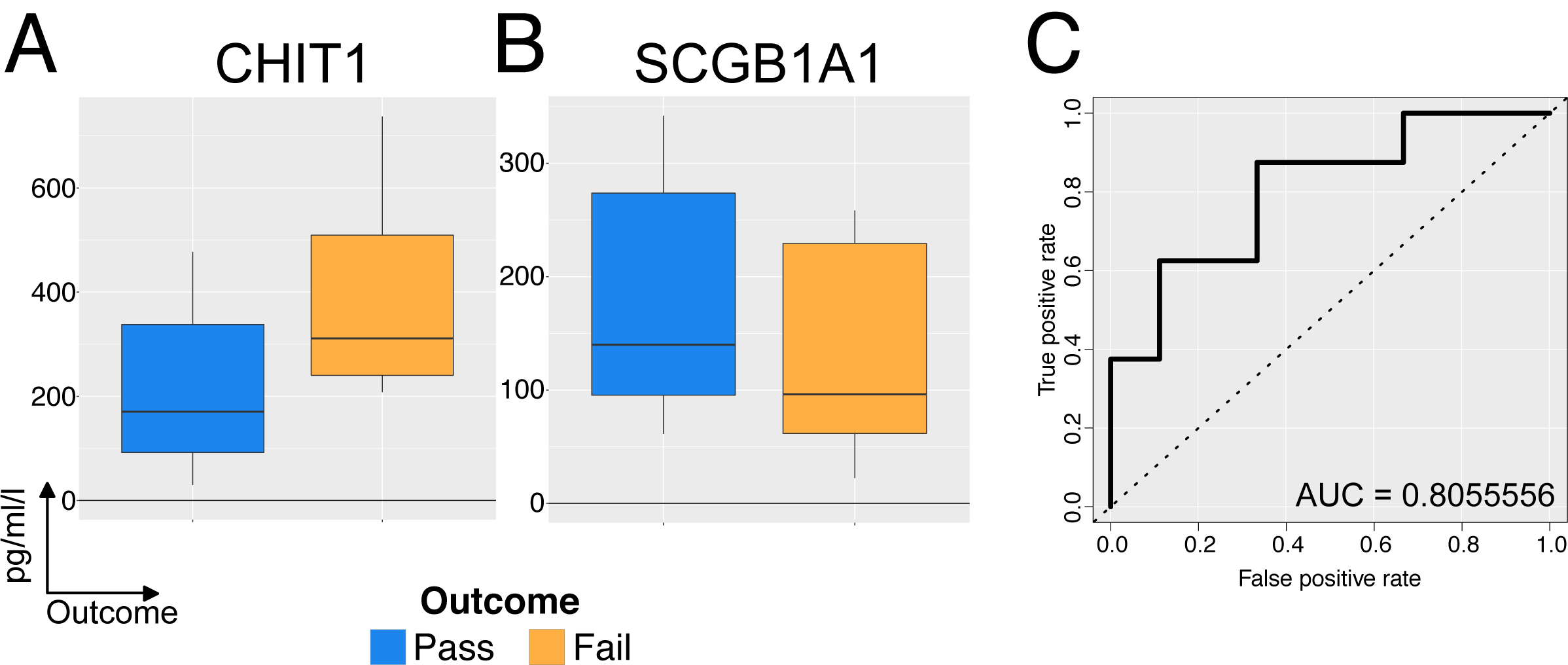
Validation of potential protein level biomarkers for success of perfusion in n=18. **A** Concentration of CHIT1 in perfusate samples taken after 150 min of perfusion with the Lund protocol. Data has been normalized for total lung capacity. **B** – Concentration of SCGB1A1 as for A. **C** A ratio of the proteins from A and B was calculated to produce a test statistic which was used for subsequent ROC analysis.

## Discussion

In order for EVLP is to be effectively utilised in clinical practice as a platform for the functional assessment, reconditioning and potential treatment of donor lungs, an increased understanding of the cellular and molecular events occurring is required. In this study, we aimed to address this knowledge gap by performing an unbiased analysis of the lung transcriptome prior to and following EVLP. This revealed that several hundred genes were differentially expressed during EVLP, with an induction in a number of innate immune (including *TNF* and *IL1B*) and HSP genes. These data suggest that the tissue damage that occurs during donor lung procurement and hypothermic transport has a substantial capacity to stimulate sterile inflammation during EVLP. Furthermore, the increase in HSPs, (which form part of a canonical cellular stress response), demonstrate the activation of cell-intrinsic compensatory mechanisms to promote organ viability. HSPs form an important part of the cellular protein quality control system that ensures misfolded or damaged proteins are degraded to prevent the initiation of apoptosis. Ischaemia reperfusion injury is a potent protein-damaging stimulus and the protein quality control machinery critical for ensuring cellular survival in this context. HSPs were originally categorised into several families according to their molecular weight, with HSPB family members among those classified as ‘small’ HSPs. Some HSPs are constitutively expressed, whilst others are induced by cell stress. Our analysis showed three patterns of HSP gene expression in lungs, and unusable lungs failed to upregulate a subset of HSP genes (dominated by HSPB genes that encode small HSPs). Of note, transgenic over-expression of the small HSP, HSPB1 (HSP27) has been shown to protect murine hearts and livers from apoptosis in the context of ischaemia re-perfusion injury (48) (49) and HSPB1 also has anti-inflammatory effects in airways epithelial cells (50). These intriguing data suggest that detection of upregulation of this HSP gene subset might be a useful indicator of more suitable lungs that warrants assessment in larger datasets.

The lungs are highly susceptible to acute injury in the critical care environment, and in the hours or days leading up to organ donation may be exposed to the sequelae of brain-stem death as well as infection, aspiration, barotrauma, fluid overload or multiple transfusions (51). The extent of donor lung injury is difficult to assess at the time of organ procurement and current donor acceptance criteria are poor discriminators of injury (3). Robust biomarkers that stratify donor lung injury and response to EVLP would assist in the rationale assessment of lungs and in the application of targeted interventions. To this end, we compared the transcriptome of lungs deemed transplantable with those that were deemed unusable. We found that the expression of immune pathway genes, including NLRP3 inflammasome-associated genes, positively correlated with lungs deemed unusable in post-EVLP biopsies. This suggests that an increased response to DAMPs, which might either be genetically determined, or may occur as a result of increased release of DAMPs due to tissue damage, is associated with worse performance during EVLP using standard physiological assessment parameters. These data mirror those obtained from candidate biomarkers studies that have associated high perfusion IL1β levels early during EVLP with worse outcomes in post-EVLP use of organs, and increased expression of cell adhesion molecules and leucocyte infiltration (19).

Our analysis of post EVLP biopsies identified 12 genes that were statistically significantly upregulated in unusable lungs, and 2 genes that were upregulated in transplantable lungs. These genes may have potential as novel transcriptional biomarkers but are also likely to reflect functionally significant pathways activated in Pass and Fail lungs. *CHIT1* (encoding chitotriosidase) was the gene most significantly upregulated in unusable lungs. Chitinase is a hydrolase expressed in plants that digests the cell walls of chitin-containing eukaryotic pathogens, such as fungi, and plays a role in pathogen defence (52). Chitotriosidase is one of two chitinases present in mammals and the most highly expressed chitinase in humans (33). Murine models suggest that CHIT1 may enhance TGFβ receptor signaling following lung injury (53). Furthermore, in humans, increased *CHIT1* expression has been found in areas of the lung affected by granulomata and fibrosis in tuberculosis, sarcoidosis, idiopathic pulmonary fibrosis, scleroderma, and chronic obstructive lung diseases. CHIT1 concentration in serum also correlates with disease progression and severity in some of these disorders, leading to the suggestion that serum CHIT1 may be a useful biomarker (33). Our data raise the possibility that CHIT1 pre-transplant expression in lungs may be a useful biomarker that identifies donor lungs likely to perform poorly during EVLP.

The chitinase-like protein *CHI3L1* was also among the significantly upregulated genes in unusable lungs. Chitinase-like proteins are evolutionarily conserved in mammals but do not have the enzymatic activity to directly degrade chitin. Rather, they appear to have evolved to play a role in the development and progression of Th2 immune responses and in defence against parasitic infections and cancer (54–56). Chitinase-like proteins are expressed by macrophages and T cells and are involved in lung inflammation, remodelling and fibrosis (35, 36) and *CHIT1* and *CHI3L1* have previously been shown to be induced in alveolar macrophages during mycobacterial infection (30). Other genes upregulated in discarded lungs included *SLC7A5* and *GBP5*. SLC7A5 is an amino acid transporter that mediates lysine influx and contributes to IL1β production via mTOR complex 1 (mTORC1)-induced glycolytic reprograming of activated human monocytes (37) whilst GBP5 promotes selective NLRP3 inflammasome assembly in macrophages in response to soluble stimuli (38). Taken together, our data suggest that the activation of alveolar/pulmonary macrophages is more marked in unusable lungs compared with those deemed transplantable. Whether this is due to higher levels of tissue damage and associated DAMPs, or due to a higher macrophage-intrinsic capacity to respond to DAMPs remains to be determined.

Two genes were significantly upregulated in transplantable lungs, MTRNR2L1, (encoding humanin-like 1), and SCGB1A1, a secretoglobin expressed by club cells (39). The humanin gene family encodes for highly similar small secretory 21-24 amino acid peptides (57) that have been shown to be protect cells from oxidative stress, starvation and hypoxia (41) and MTRNR2L1 transcription is increased by oxidative stress and toll-like receptor 4 agonists (58), therefore, its upregulation may enhance cellular integrity in lungs in the face of cell stressors associated with the organ procurement process. Club cells are the major secretory cells of the human small airways. A recent single cell RNA sequencing analysis of small airways cells identified SCGB1A1 (also known as uteroglobin or CC10) as a canonical marker of club cells, and cell trajectory analysis indicate that they may have the capacity to both de-differentiate into basal cells (with stem capacity) and trans-differentiate into ciliated epithelial cells. Analysis of cell-specific gene expression suggests that club cells may also play novel roles in host defence and have anti-protease activity via the expression of α1-antitrypsin (an important neutrophil elastase inhibitor) and SLPI (secretory leucoprotease inhibitor, a potent serine protease inhibitor) (39). SCGB1A1 has been shown to have direct anti-inflammatory actions inhibiting NFkB activation and CXCL8 secretion in airway epithelial cells(42) (43). Our data raises the possibility that these cells function to protect lungs from some of the deleterious processes occurring during the organ donation and procurement process and to inhibit collateral damage caused by neutrophils recruited following reperfusion. It will be possible to test this hypothesis in future studies by investigating club cell number and activation in pre-implantation biopsies in a larger cohort of transplanted lungs.

The major limitation of our study is the relatively small sample size (n=10) but this is not dissimilar to other transcriptomic studies in lung transplantation (59, 60). However, our use of paired pre- and post-EVLP samples, as well as gene pathway analysis, substantially increased our statistical power to demonstrate meaningful increases in functionally relevant molecular pathways. Our study identified a number of novel RNA biomarkers that may indicate successful EVLP and we went on to measure the protein concentration of two of these biomarkers, CHIT1 and SCGB1A1, in perfusate in a larger validation cohort. This analysis indicated that these two measurements can be combined to predict successful EVLP with reasonable sensitivity and specificity, and merit prospective assessment in future clinical trials. Our results also suggest a number of potential targets for therapeutic intervention to improve lung suitability during EVLP that will require validation in larger studies. In particular, inhibition of the NLRP3 inflammasome and enhancement of club cell function may be of utility. The former is readily translatable, given the development of these agents for the treatment of sterile inflammation in other contexts, such as crystal-induced arthropathies and autoinflammatory conditions (61).

In summary, our data suggest that lungs deemed suitable for transplantation following EVLP have reduced induction of multiple immune pathways and are better able to generate ATP. Our study identifies specific biomarkers that may hold utility in identifying suitable lungs and pathways amenable to therapeutic intervention during EVLP that will inform the design of future clinical trials in this area.

## Methods

### Study subjects and protocol

In this study, we utilised samples of whole donor lung tissue collected from a large cohort of highly characterised clinical EVLP procedures performed as part of the DEVELOP-UK multicentre trial, which involved all five UK transplant centres(26). DEVELOP-UK included 53 adult donor lungs which were deemed unsuitable for lung transplantation but met pre-defined criteria for EVLP. Assessments were performed using a Vivoline LS1 EVLP circuit (Vivoline Medical AB, Lund, Sweden) following 1 of 2 standardized perfusion protocols: an initial Hybrid protocol featured an open left atrium, acellular perfusate, and perfusate flow limited to 40%– 60% of donor calculated cardiac output *(n=22)*. Subsequently, the perfusion strategy was changed to the Lund protocol with cellular perfusate (haematocrit 10%–15%) and full flow perfusion of 100% donor calculated cardiac output *(n=31)*. Assessment methods, including sampling procedures, remained unchanged between the two protocols. A full report of the outcomes of the DEVELOP-UK study have been previously published (26).

### Sample collection

A standardised sampling protocol was followed to allow for tissue biopsy collection from either the right middle lobe or lingula both before and after the EVLP assessment using a GIA surgical stapler(26). Biopsies were snap frozen in liquid nitrogen as soon as possible after collection for subsequent RNA isolation.

A control perfusate sample was collected from the primed EVLP circuit before donor lung perfusion started. Repeated perfusate samples (5 ml) were then collected at 15 and 30 minutes after perfusion commenced and every 30 minutes thereafter. The perfusate samples were centrifuged at 180g for 6 minutes at 4°C to remove cellular debris and then aliquoted into tubes before being frozen initially at −20°C and then transferred to a −80°C for longer term storage and subsequent laboratory analysis.

### RNA extraction

RNA was extracted from small pieces of tissue snap frozen at the indicated time points. Samples were lysed in Lysis buffer (Invitrogen) using a precellys homogenizer (Bertin Instruments), RNA extracted using a Pure link RNA mini kit (Invitrogen) and contaminating DNA removed using Turbo DNase kit (Invitrogen). RNA was quantified using by absorbance and 280nm (Nanodrop spectrophotometer) and quality assessed using a 2100 bioanalyzer (Agilent) with a RNA nano chip (Agilent).

### RNASeq

RNAseq Libraries were made using TruSeq Stranded Total RNA library prep kit (Illumina) with 7 minutes fragmentation as per manufacturers instructions. Final libraries were amplified for 14 cycles by PCR and the final size assessed using a 2100 bioanalyzer (Agilent) with a HS DNA chip (Agilent). Pooling of finally libraries and sequencing was carried out by Eurofins on a Hiseq 2500 on a Rapid run 1×50.

### RNASeq analysis

BCL files were demultiplexed using CASAVA and resulting fastq files aligned to the hg38 genome using hisat2. Gene level counts were determined using Featurecounts from RSubread and a count table produced. Differential expression analysis was carried out using DESeq2 after fitting an appropriate model within the R statistical environment.

### scRNAseq analysis

The dataset GSE103354 was used to produce a club cell specific gene signature (44). Briefly the publicly available data was analysed using the standard Seurat pipeline. Following normalisation and scaling of the data clusters were identified and annotated. Marker genes for the club cell cluster were determined and the top 100 as ranked by log fold change were used as the Club cell specific gene set for subsequent analysis.

### Gene Set Enrichment Analysis

For GSEA comparisons of interest were ranked by the inverse of the raw p value with the sign of the log fold change such that the gene ranked 1 showed the most significant positive fold change and the nth gene the most significant negative fold change. Genes which showed no difference are centred around nth/2 rank. GSEA was subsequently carried out using the java based GSEA applet maintained by the Broad using the classic enrichment statistic in pre-ranked mode.

### Receiver Operating Characteristic Analysis

A test statistic for prediction of EVLP outcome was produced by taking the ratio of the concentration of CHIT1 to SCGB1A1 protein present in perfusate at 150 min post perfusion from 17 perfused lungs made up from the DEVELOP UK study including 7 from the set of samples used for the RNA-seq analysis. The R packaged ROCR was used to perform the test and an area under the curve was calculated.

### MSD methods

All protein expression measured in perfusate was adjusted to the predicted total lung capacity (pTLC) of the donor as an estimate of perfused donor lung volume and were reported as corrected perfusate concentrations (pg/ml). The pTLC was calculated in a routine fashion based on donor gender and height. Interleukin (IL)-1b, IL-6, IL-8, TNF-a, and MCP-1 were analysed with an MSD Multi-ArrayVR (Meso Scale Diagnostics, LLC, Rockville, MD, USA). The assay was performed according to the manufacturer’s instructions.

### Enzyme-linked immunosorbent assays

Serial perfusate samples from 17 human donor lungs undergoing clinical EVLP assessments were analyzed retrospectively. Enzyme Linked Immunosorbent Assays (ELISA) were used to detect protein levels in the perfusate for CHIT1 (BioTeche Novus Biologicals, Abingdon, Oxfordshire, UK), SCGB1A1 and soluble Intercellular Adhesion Molecule 1 (ICAM-1) (R&D Systems, Inc., Minneapolis, MN, USA). All protein concentrations measured in perfusate were adjusted to the donor predicted total lung capacity as an estimate of perfused donor lung volume and reported as corrected perfusate concentrations (pg/ml).

### Figure generation

Some of the illustrations contain images produced using ©BioRender.

## Data availability

Original Data will be made available upon all reasonable requests to the corresponding author.

## Acknowledgements

This work was funded by the National Institute for Health Research Blood and Transplant Research Unit (NIHR BTRU) in Organ Donation and Transplantation at the University of Cambridge and Newcastle University and in partnership with NHS Blood and Transplant (NHSBT). The views expressed are those of the author(s) and not necessarily those of the NHS, the NIHR, the Department of Health or NHSBT.

J.R.F., C.C and M.M are supported by the NIHR BTRU.

M.R.C. is supported by the NIHR Cambridge Biomedical Research Centre, a Medical Research Council New Investigator Research Grant (MR/N024907/1), and an NIHR Research Professorship (RP-2017-08-ST2-002).

## Author Contributions

JRF, MM, AF and MRC conceived and designed the research project. JRF, MM, CC, AC and LEB carried out experiments. AA and WES collected, processed and stored samples. JRF analysed the data. JRF and MRC interpreted the data. MM, JRF and MRC wrote the manuscript, and AF edited the manuscript. All authors reviewed and commented on the manuscript.

---

**Supplemental Figure 1.**
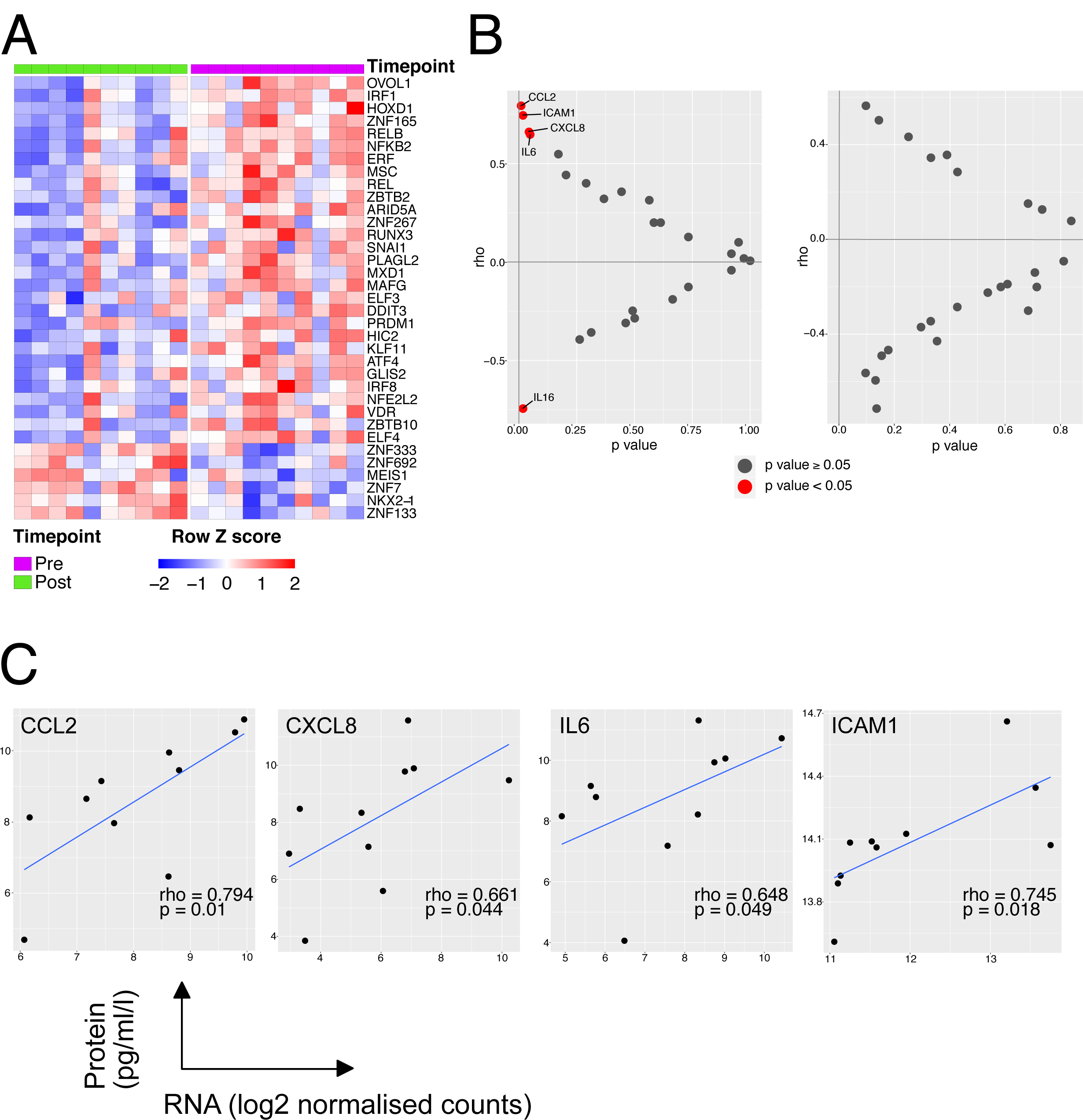
**A** – Heatmap of all genes identified as transcription factors in the TFCheckpoint database and significantly differentially expressed during EVLP **B** - Correlation of transcriptome either pre (Left) or post (Right) EVLP with cytokines in the perfusate at the end of the procedure. Cytokines were measured in perfusate using a multiplex MSD kit. The normalised RNA level in the samples prior to EVLP was correlated using Spearmans method with the protein level detected in the perfusate. Correlations with a p value of less than 0.05 have been labelled. **C**-Scatter plot of normalised transcript expression prior to EVNP against protein level in the perfusate at the end of EVLP. Correlation coefficient and p values was calculated using Spearmans test prior to log2 transformation, blue line indicated a liner model fitted to the data post transformation. Cytokines compared are indicated at the top of the respective plot.

**Supplemental Figure 2.**
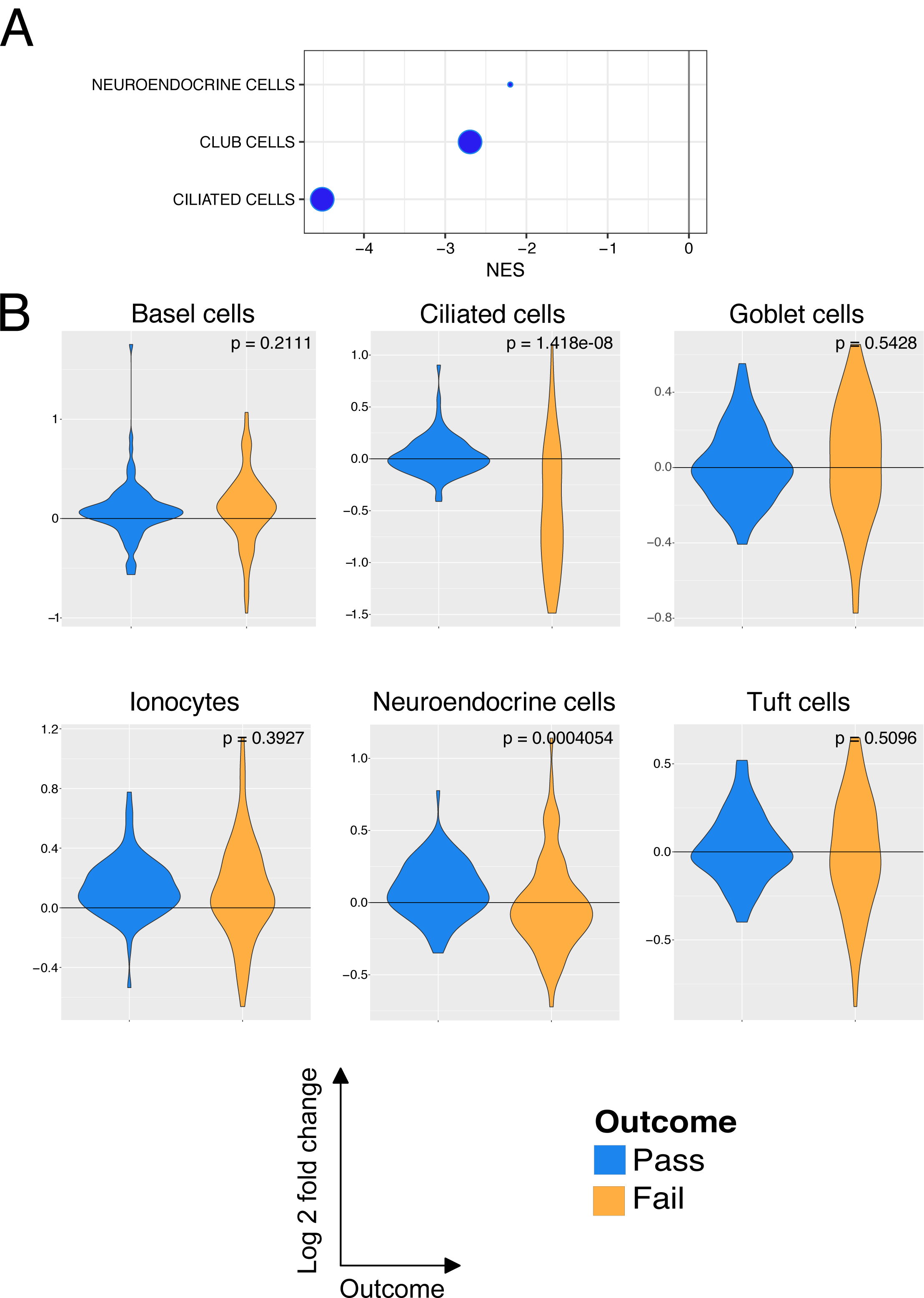
**A** – GSEA analysis for all lung cell subset marker genes following EVLP comparing those lungs which passed and those that failed EVLP. All significantly enriched pathways have been plotted, size of point is inversely correlated to the FDR q value, red points indicated positively enriched pathways and blue negatively enriched. **B** - Violin plot indicated the log fold change in expression for all lung cell subset marker genes. The p value is for comparison of the two groups using a Mann-Whitney test. Blue is organs which have passed EVLP and orange genes which have failed.

**Supplemental Figure 3.**
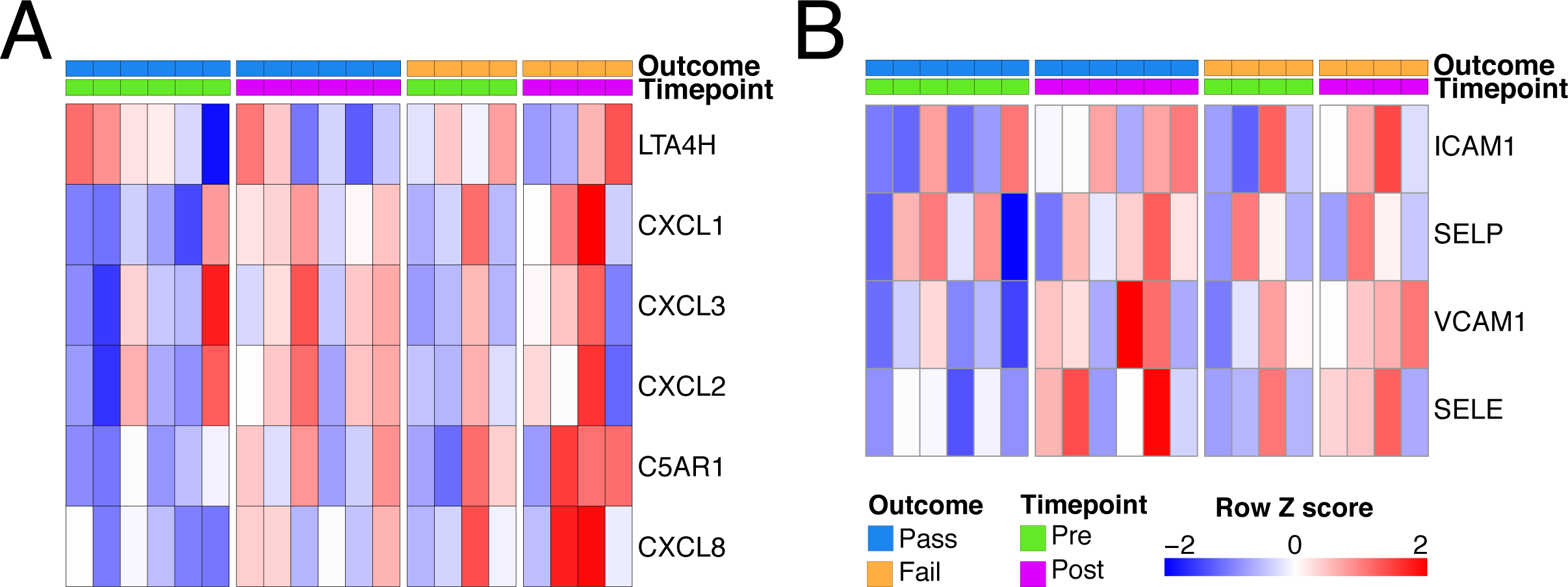
Neutrophil modulation during EVLP. **A** - Heatmap of genes involved in neutrophil chemoattraction. **B** - Heatmap of genes involved in neutrophil adhesion.

